# Perivascular RELM⍺-positive synovial macrophages recruit monocytes at the onset of inflammatory arthritis

**DOI:** 10.1101/2025.02.11.637672

**Authors:** Barbora Schonfeldova, Marah Chibwana, Rebecca Gentek, Kristina Zec, Irina A Udalova

**Author notes:** Contributed equally (co-last authors) (Kristina Zec, Irina A Udalova).

## Abstract

Macrophages, monocytes and neutrophils are major types of myeloid cells involved in inflammatory diseases, such as rheumatoid arthritis (RA). Recent scRNA-seq studies identified a remarkable diversity of synovial macrophages but, with the exception of lining macrophages, their geographical location and specific roles remain largely unexplored. Here, we localised the RELM⍺-positive macrophages, predicted to produce high levels of monocyte-recruiting chemokines, to the synovial interstitium and more specifically, to the vicinity of interstitial blood vessels. Using complementary reporter mouse models, CCL2^mCherry^ to label CCL2-producing cells, and CCR2^CRE/mKate2^ marking CCR2 expressing monocytes, we demonstrated that RELM⍺-positive perivascular macrophages secrete CCL2 to recruit monocytes predominantly to the synovial interstitium at the onset of antigen-induced arthritis. The inflamed synovial environment guides the differentiation of the recruited monocytes into tissue-resident macrophages, including but not limited to macrophages expressing VSIG4, a characteristic marker of lining macrophages. Thus, RELMα-positive macrophages orchestrate monocyte recruitment to the synovium during articular inflammation, contributing to a local replenishment of synovial lining macrophages.

## Introduction

The synovium is a soft tissue that lines the diarthrodial joints, tendons, and bursae [1]. Improvements in synovial sampling and spatiotemporal profiling on protein and RNA levels greatly enhanced our understanding of this tissue site. It is now understood that the synovium in both mice and humans is highly heterogeneous both at steady-state and during inflammatory diseases [2-6]. Among important players in the synovium are the synovial macrophages, whose markers were previously identified by single-cell RNA sequencing [3]. Across multiple tissues and species, interstitial macrophages were previously identified to be broadly categorised by localisation as either vasculature-associated or innervation-associated [7]. However, the localisation of interstitial synovial macrophages identified as RELM⍺-positive and MHCII-positive macrophages in the synovium has not been previously studied. Similarly, whether either of these subsets has a role in the recruitment of immune cells, including monocytes, during inflammatory arthritis is unknown. Therefore, we decided to profile the RELM⍺-positive synovial macrophages and identify their localisation and function during inflammatory arthritis.

In this brief research report, we identified that RELM⍺+ macrophages are located in the perivascular niche and are the principal producers of CCL2, and guide monocytes recruitment to the interstitium at the onset of synovial inflammation. The recruitment of monocytes is an important mechanism both during steady-state and during inflammation to (1) promote the recruitment of other immune cells and (2) replenish local tissue-resident subsets of macrophages. Dissecting the local signals and mechanisms leading to adaptation of different macrophage differentiation trajectories by monocytes will further our understanding of the role these cells play during inflammatory arthritis.

## Methods

### Animals

Mice were maintained in specific pathogen-free conditions in the Kennedy Institute of Rheumatology under the establishment licence of the University of Oxford. All animal work complied with the Animal (Scientific Procedures) Act 1986, following the national and institutional standards.

Animal strains used: CCR2^CreER mKate2^ mice [8] were used to trace monocyte localisation, CCL2^mCherry^ strain (JAX ID: 016849) [9] was used to locate CCL2-producing cells, Ms4a3^tdTomato^ Cx3cr1^eGFP^ [10] strain was used for tracing monocyte contribution to lining macrophage replenishment at day 28 of AIA, and C57BL/6 mice were used for all other experiments.

### *In vivo* model of antigen-induced arthritis

The induction of antigen-induced arthritis (AIA) was performed in a modified version of a previously described protocol [11]. Firstly, mice were immunised by two subcutaneous injections with an emulsion of methylated bovine serum albumin (mBSA) (at a concentration of 40mg/ml, Merck, Cat# A1009) and complete Freund’s adjuvant (3.3mg/ml, Scientific Labs, Cat# 263910) suspended in PBS. The emulsion was created using the BTB immunisation kit (BTB Emulsions, Cat# PK-M-1-NP). A standardised method using a shaking homogeniser to prepare adjuvant/ antigen emulsions to induce autoimmune disease models was used [12, 13]. Emulsions were prepared according to the manufacturer’s recommendations. After a week, mice were challenged locally by intra-articular injection of either mBSA (20mg/ml, left knee) or PBS (right knee, control) by Hamilton 50 μl syringe with Luer tip. Both subcutaneous and intra-articular injections were performed under isoflurane anaesthesia.

### Preparation of murine knee sections

Murine knee section preparation was done as previously described [14]. Essentially, the knees of naïve or mice with AIA, sciatic neurectomy or PMX were isolated and fixed for 4 hours at 4°C in 4% methanol-free periodate-lysine-paraformaldehyde (PLP). Samples were then washed 3 times in PBS and moved into a decalcification buffer (distilled water, 0.5M EDTA, pH 7.4) for a week. The decalcification buffer was refreshed once during this time. Samples were then moved to 30% sucrose for cryoprotection for 24 hours until they sank. Then, they were embedded in an embedding medium (8 g of gelatine, 2 g of PVP, 20 g of sucrose in 100 mL of PBS). The embedded tissue was left at room temperature (RT) until the embedding medium solidified and then frozen in methanol and dried ice slurry. The tissue blocks were then stored at -80°C until they were sectioned at 20μm thickness at Leica cryostat CM1900 and mounted onto gelatine-coated glass slides. Those were subsequently dried for 1-2 hours at RT and then kept at -20°C.

### Immunofluorescent labelling of murine knee sections

Knee sections were dried at RT for 40 minutes and at 65°C for an additional 20 minutes. Sections were rehydrated with PBT (PBS, 0.05% Triton X-100) for 5 minutes at RT in a humidified chamber. The sections were subsequently incubated with Image-iT™ FX Signal Enhancer (Invitrogen, cat# R37107) for 30 minutes at RT. Slides were briefly rinsed in PBS and 1X Carbo-free blocking buffer (deionised water, 10X Carbo-free (Vector Laboratories, Cat# SP-5040), 0.3M glycine, 0.08% NaN_3_, 0.05% Triton X-100) was applied for at least 1 hour at RT in a humidified chamber. Sections were then stained by primary antibodies in the 1X Carbo-free blocking buffer at concentration 1μg/slide overnight at 4°C in a humidified chamber. The next day, slides were washed 3 times in PBS for 5 minutes at RT. Sections were then stained with secondary antibodies (concentration 2μg/slide) diluted in 1X Carbo-free blocking buffer for 1.5 hours at RT in a humidified chamber. Slides were then washed 3 times in PBS for 5 minutes at RT and incubated with 2μM of Sytox blue dye (Invitrogen, Cat# S34857) diluted in PBT for 25 minutes at RT. Slides were quickly immersed in deionised water and subsequently mounted using FluorSave™ (Merck, cat# 345789) and coverslips. Slides were then dried overnight at RT in the dark and imaged within a week using a Zeiss980 confocal microscope with a 20X objective.

### Synovial single-cell suspension for flow cytometry

Synovial isolation was performed using a modified version of a previously published protocol [15], described in detail in Zec et al 2023 [16]. Isolated synovium was digested with DNase I (100μg/ml, Merck, Cat# 11284932001) and Liberase TL (0.4mg/ml, Merck, Cat# 540102001) in RPMI 1640 (1% Pen-Strep) for 1 hour at 37°C, shaking at 200rpm. The remaining tissue was crushed through a 100 μm cell strainer and washed with RPMI 1640 (1% Pen-Strep) medium.

### Flow cytometry staining

Synovium single-cell suspensions were stained with near-IR live/dead dye (1:500, Invitrogen, Cat# L10119) in PBS for 15 minutes at 4°C. Then, the synovium samples were blocked with Fc block (BD Biosciences, Cat# 553142) diluted 1/100 in FACS buffer for 10 minutes at 4°C. Primary antibodies were diluted in 1X Brilliant stain buffer (BD Biosciences, Cat# 563794) in FACS buffer and cells were stained with them for 30 minutes at 4°C. Cells were fixed for 10 minutes at 4°C with 25μL of Cytofix (BD Biosciences, Cat# 554655). Cells were permeabilised with 1X Perm/Wash buffer (BD Biosciences, Cat# 554723) for 20 minutes at 4°C. RELM⍺ and CCL2 were stained intracellularly with 1X Perm/Wash buffer for 30 minutes at 4°C. All antibodies and dilutions used for flow cytometry are summarised in Supplementary Table 1.

**Supplementary Table 1:** Primary and secondary antibodies used.

| Epitope | Colour | Clone | Manufacturer | Cat # | Dilution used |
| --- | --- | --- | --- | --- | --- |
| CCL2 | PE | 2H5 | Biolegend | 505903 | 1/50 |
| CD11b | BV785 | M1/70 | Biolegend | 101243 | 1/200 |
| CD31 | BV421 | MEC 13.3 | BD Biosciences | 562939 | 1/80 |
| CD45 | BV650 | 30-F11 | Biolegend | 103151 | 1/100 |
| CD68 | Unconjugated | FA-11 | Biolegend | 137001 | 1/200 |
| F4/80 | PE/Dazzle™ 594 | BM8 | Invitrogen | 123145 | 1/200 |
| Live/Dead | Near-IR | N/A | Invitrogen | L10119 | 1/500 |
| Ly6C | BV711 | HK1.4 | Biolegend | 128012 | 1/100 |
| Ly6G | FITC | 1A8 | Biolegend | 127605 | 1/100 |
| mCherry | AF594 | 16D7 | Invitrogen | M11240 | 1/200 |
| MHCII | AF700 | M5/114.15.2 | Biolegend | 107622 | 1/100 |
| MHCII | BV421 | M5/114.15.2 | Invitrogen | 404-5321-82 | 1/100 |
| RELMα | PerCP-eF710 | DS8RELM | Invitrogen | 46-5441-82 | 1/100 |
| RELMα | Unconjugated | Polyclonal | Abcam | ab39626 | 1/80 |
| RFP | Unconjugated | Polyclonal | Invitrogen | R10367 | 1/400 |
| VSIG4 | APC | NLA14 | Invitrogen | 17-5752-82 | 1/100 |
| VSIG4 | Unconjugated | Polyclonal | Bio-Techne (R&D Systems) | AF4674 | 1/80 |
| Secondary antibodies |  |  |  |  |  |
| Colour | Species reactivity | Manufacturer | Cat # | Dilution used |  |
| BV421 | Goat | Strattech Scientific Ltd | 705-675-147 | 1/20 |  |
| AF488 | Rat | Invitrogen | A-21208 | 1/400 |  |
| AF594 | Rabbit | Invitrogen | A-21207 | 1/400 |  |
| AF647 | Rabbit | Invitrogen | A32795 | 1/400 |  |

### Single-cell RNA sequencing analysis

The publicly available scRNAseq dataset profiling murine myeloid cells in the synovium [3] was accessed under the accession code GSE134691 from Gene Expression Omnibus. The Seurat V3 package [17] was used to re-analyse the day 1 dataset. A quality control check was performed with a cut-off for mitochondrial genes 6 and an RNA count of 14,000. The dimensionality of the dataset was investigated, and 22 principal components were used for subsequent analysis. The clustering was assessed by the clustree package [18] and clustering resolution 0.5 was selected to determine the different subsets of macrophages. The Gsfisher package, developed by Prof Steve Sansom and Dr Kevin Rue-Albrecht, was used to assess the gene ontology of synovial macrophages.

## Results

### 3.1 RELM⍺-positive macrophages are localized in the vicinity of interstitial vasculature

A recent comprehensive scRNAseq analysis of synovial myeloid cells recognised the interstitial macrophages as expressing either major histocompatibility complex II (MHCII) or Resistin-like molecule alpha (RELM⍺) [3]. Using immunofluorescent (IF) staining of naïve murine knee synovial sections, with antibodies to MHCII, RELM⍺, and a pan-macrophage marker IBA1, we examined whether these markers mark distinct macrophage subsets. Sections were collected from CX3CR1^eGFP^ mice and the CX3CR1 signal was used to mark lining macrophages at a steady state, as previously described [3, 16, 19]. We noted that whilst some interstitial macrophages co-expressed MHCII and RELM⍺, many macrophages in the interstitium expressed only one of the markers (**Supplementary Figure 1**). Thus, we hypothesised the two subsets of interstitial macrophages may have distinct localisation and function.

Since little is known about RELM⍺-positive macrophages in the synovium, we decided to further investigate this population. RELM⍺-positive macrophages in other organs, were localised either in the vicinity of the vasculature or innervation [7]. However, in the human synovium, the counterpart of RELM⍺-positive macrophages was identified to express high levels of LYVE1, suggesting their possible perivascular localisation [4, 7]. To test this hypothesis, we utilised fluorescence confocal microscopy and co-stained murine knee sections with a marker of vasculature CD31, a marker of innervation tubulin β3 chain (Tubb3), a pan-macrophage marker CD68 and RELM⍺.

We noted that RELMα-positive macrophages were localised in the interstitium, often in the proximity to CD31-positive vasculature, rather than the innervation (**Figure 1A-D**). Interestingly, the larger vessels were often associated with innervation, whereas most RELMα-positive macrophages we observed closer to the smaller vessels, devoid of innervation (**Figure 1B**). Conversely, CD68-positive macrophages localized close to the Tubb3-positive innervation were negative for RELMα (**Figure 1C**). The quantification of the confocal microscopy images confirmed the preferential localisation of RELM⍺-positive macrophages in the vicinity of blood vessels rather than Tubb3-positive innervation (**Figure 1E, F**).

**Figure 1:**
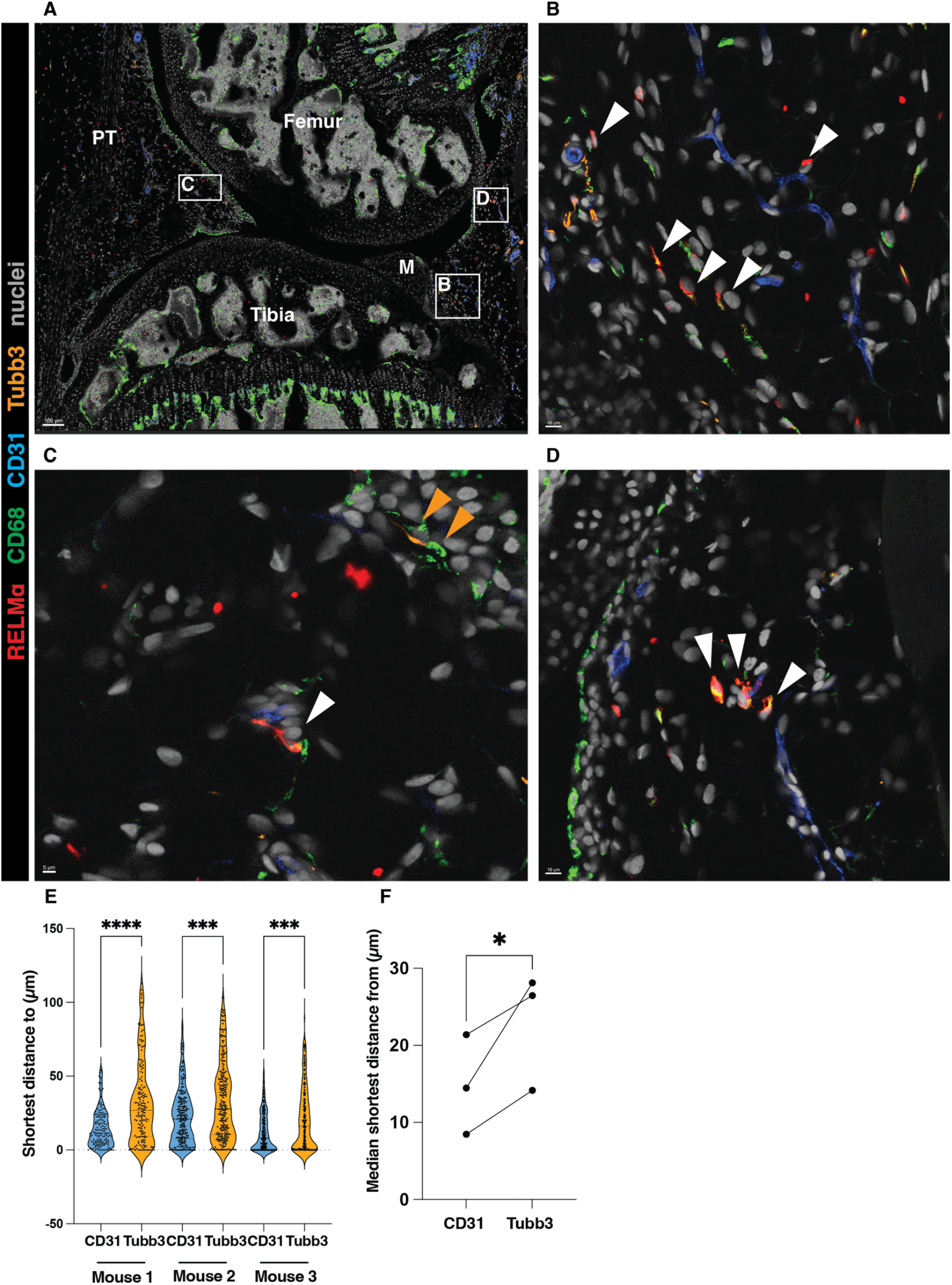
RELM⍺-positive macrophages are localised in the vicinity of synovial blood vessels. **(A-D)** A representative image showing the localisation of RELM⍺-positive macrophages in the vicinity of the synovial vasculature (marked by CD31). The white boxes represent ROIs enlarged in B, C, and D. White arrows indicate RELMα-positive macrophages in the vicinity of the vasculature; orange arrows show CD68-positive macrophages close to the Tubb3-positive innervation. M=meniscus, PT=patellar tendon. **(E-F)** Quantification of the localisation of RELM⍺-positive macrophages in respect to the vasculature (CD31) or innervation (Tubb3), at least 2 positional duplicates/mouse were quantified. Raw data shown in (E) were analysed by the non-parametric Kruskal-Wallis test with Dunn’s multiple comparison test; the median of the positional duplicates/mouse shown in (F) were compared by parametric paired t-test.

### 3.2 RELM⍺-positive macrophages produce CCL2 during antigen-induced arthritis (AIA)

The function of RELM⍺-positive macrophages in the synovium is unknown. To predict it, we used the publicly available single-cell RNA sequencing dataset from the onset of serum transfer-induced arthritis (STIA) [3]. We identified that on day 1 of STIA, RELM⍺+ macrophages were expressing genes which were part of the GO pathway: monocyte and cell chemotaxis (**Figure 2A**). More specifically, they were indicated as the main producers of the CCR2 ligands *Ccl2, Ccl7*, and *Ccl8* [20, 21] (**Figure 2B-D**) at the onset of STIA [3], suggesting that RELM⍺-positive macrophages may represent the tissue-resident macrophage subset capable of recruiting monocytes. As CCL2 is a major monocyte chemoattractant and highly important for the pathology of inflammatory arthritis [20, 22, 23], we wanted to examine whether RELM⍺+ macrophages produce CCL2 on the protein level at the onset of AIA as well (**Figure 2E**). The AIA model is a well-described model of inflammatory arthritis with distinct stages of inflammation, making it a well-suited model for studies on early myeloid cell recruitment [16, 24].

**Figure 2:**
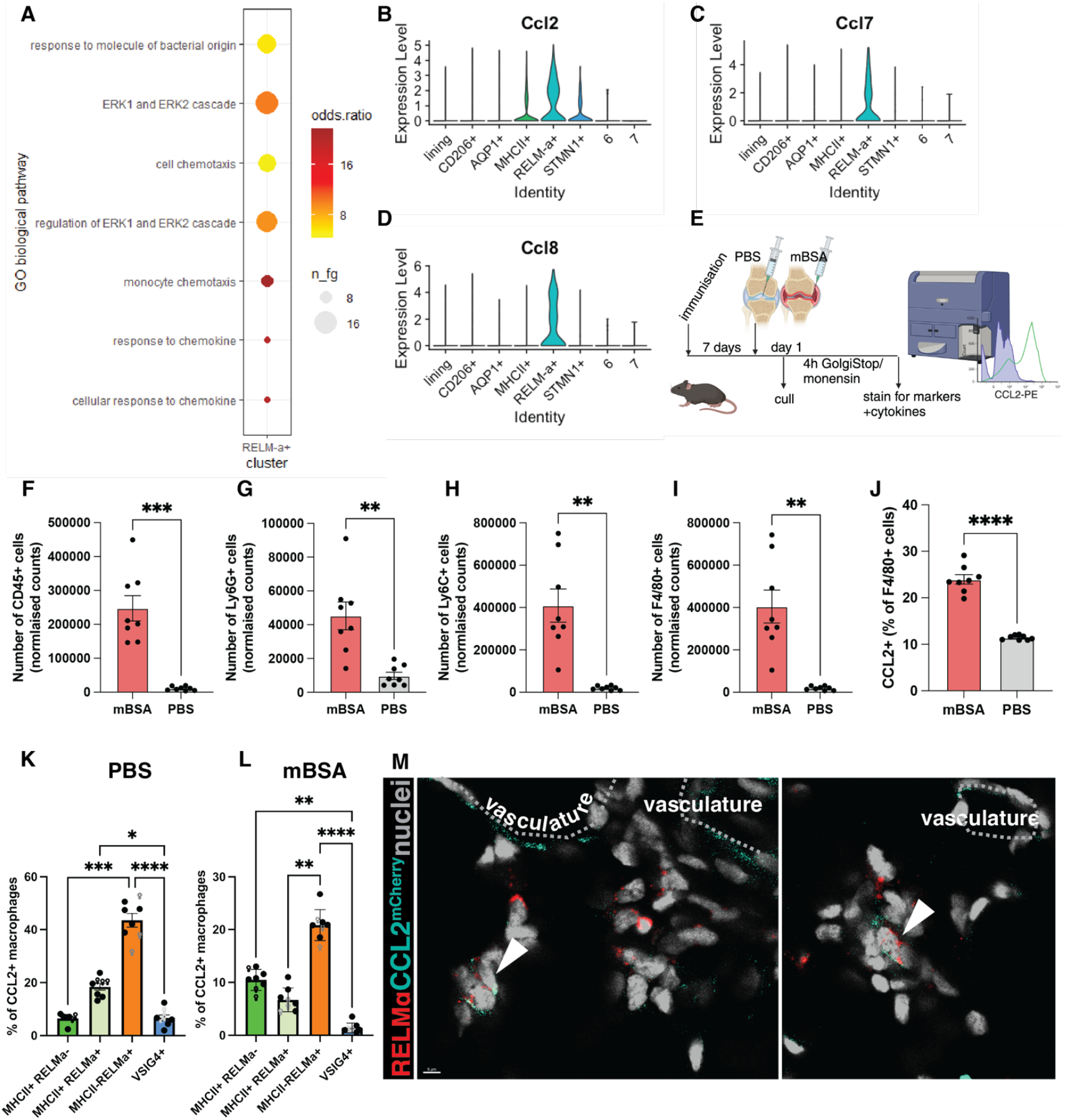
RELM⍺-positive macrophages produce CCL2 during the onset of AIA. **(A)** Predicted GO biological pathways in the RELMα-positive macrophages, re-analysis of day 1 of STIA [3] using gsfisher. Dot plot showing the enrichment of the most significant categories in a subset of interest. n_fg = number of genes in the category, odds ratio = effect size (over-representation). **(B-D)** VlnPlot showing the expression of *Ccl2* (A), *Ccl7* (C), and *Ccl8* (D) from day of STIA [3]. **(E)** The experimental plan for assessing CCL2 production at the onset of AIA. **(F-I))** Normalised counts of leukocytes (F), monocytes (G), neutrophils (H), and macrophages (I) at day 1 of AIA in the antigen-injected knee (mBSA) and control (PBS). **(J)** Percentage of CCL2-producing macrophages within all macrophages in mBSA vs PBS. **(K-L)** Percentage of CCL2-producing subset of macrophages within CCL2-producing macrophages identified in (E) in PBS (F) and mBSA (G) injected knees. Each point represents a biological replicate, a total of n=8. ROUT identified outliers with Q=1%, and normality was tested by the Shapiro-Wilk test. Paired t-test (F-J) and Kruskall-Wallis with Dunn’s post hoc multiple comparison test (K-L) were used to identify significant differences between experimental groups. * < 0.05, ** < 0.01, *** < 0.001, **** < 0.0001 **(M)** Example images of RELM⍺-positive macrophages at 6 hours p.c. in the AIA model expressing CCL2; imaging was done on CCL2^mCherry^ mice; the dotted-line highlights vasculature.

CCL2 production by synovial macrophages on day 1 of AIA was assessed by flow cytometry (**Supplementary Figure 2**). As expected, on day 1 of AIA, there is an increase in the number of immune cells, such as neutrophils, monocytes, and macrophages in the antigen-injected knee compared to the contralateral PBS-injected knee (**Figure 2F-I**). Notably, the percentage of CCL2-producing macrophages was also significantly increased (**Figure 2J**). Next, we profiled which subset of tissue-resident macrophages was the primary producer of CCL2 at the onset of AIA. As anticipated from the scRNAseq re-analysis, RELM⍺-positive macrophages were the principal subset of tissue-resident macrophages producing CCL2 on the protein level (**Figure 2K, L**). Interestingly, in the antigen-injected joint, tissue-resident macrophages comprised less than half of the F4/80-positive cells producing CCL2 (**Figure 2L**), suggesting that newly infiltrated macrophages assume this function upon entering the synovium.

To localise CCL2 production at the onset of AIA, we utilised the CCL2^mCherry^ mouse model, in which the *Ccl2* 3’ end is tagged with mCherry and noted the abundance of mCherry-fused CCL2 within the interstitial RELM⍺-positive macrophages at the onset of AIA (**Figure 2M**).

### 3.3 Monocytes are recruited to the interstitium at the onset of inflammatory arthritis

Synovial lining macrophages were previously shown to recruit neutrophils to the lining niche at the onset of inflammatory arthritis via secretion of CXCL1 [16]. However, it is unknown whether monocytes follow the same localisation pattern as neutrophils or whether these early responders are recruited into a different area during synovial inflammation. To investigate monocyte recruitment during AIA, we used the CCR2^CRE/mKate2^ mouse model, where monocytes are marked by fluorescent marker mKate2 [8]. A very few CCR2+ monocytes were observed in the synovium at the steady state (**Figure 3A**), but at 6 hours post-challenge, their number dramatically increased in the interstitium near blood vessels (**Figure 3B**). At day 2 of AIA, which is the peak of inflammation in this model, the interstitial localisation of monocytes is less apparent (**Figure 3C**), as the synovium is filled with infiltrating monocytes. Similarly, neutrophils were previously reported to lose their lining niche localisation pattern at the peak of inflammation [16]. To confirm our observations of monocyte localisation, we quantified the proximity of monocytes to interstitial compared to the lining vasculature. As expected from our representative images, monocytes were strongly associated with interstitial vasculature rather than with the lining vasculature at the onset of AIA (**Figure 3D, E**), in contrast to neutrophil recruitment and localisation pattern [16, 19].

**Figure 3:**
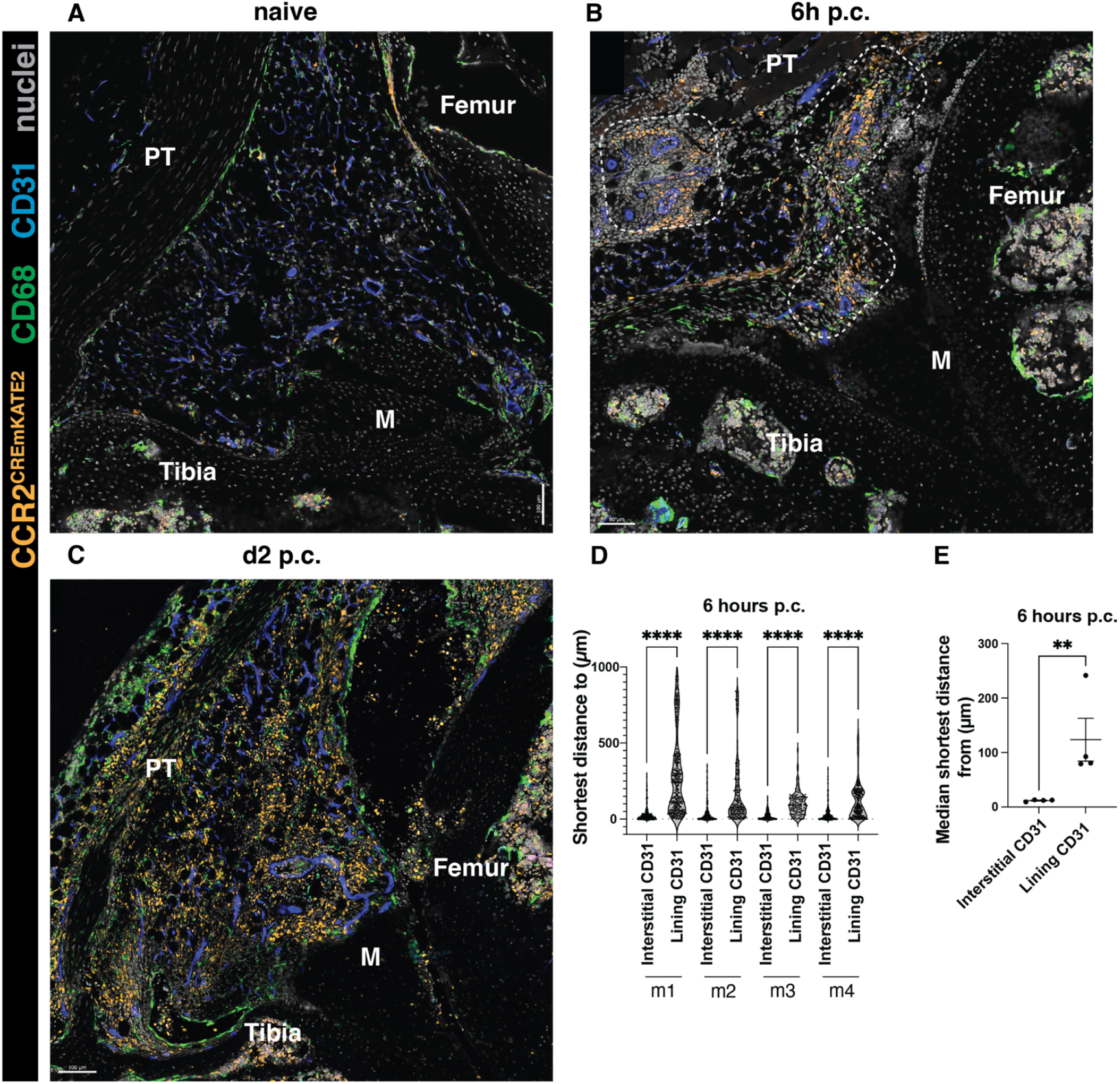
Monocytes are recruited to the interstitium at the onset of AIA. **(A-C)** Representative images of CCR2^CreERT-mKATE2^ murine knee sections at steady-state (A), and at the onset (B, 6h p.c.), and peak (C, day 2 p.c.) of AIA. **(D-E)** Quantification of the localisation of CCR2-positive monocytes at 6h post-challenge. (D) shows raw data displaying every cell, each point represents a single monocyte in a given biological replicate. In (E), each point represents a biological replicate (n=4). Kruskal-Wallis test was used to identify differences in (D) and a parametric ratio paired t-test was used to quantify differences in (E).

### 3.4 Monocytes can give rise to the lining macrophages during the resolution phase of inflammatory arthritis

During the resolution phase of the AIA model, at day 14 post-challenge, we observed that some CCR2+ monocytes remained in the synovium, but their localisation was now in proximity to the synovial lining (**Figure 4A**). This increased presence of monocytes during resolution compared to naïve knees was corroborated by flow cytometry quantification (**Figure 4B**). Notably, this phenomenon was only observed in the antigen-injected mBSA knee joint and not the contralateral PBS-injected knees (**Figure 4B**). This indicated that monocytes are retained in the synovium and could potentially replenish tissue-resident macrophages lost during inflammation. Indeed, at day 14 post-challenge, some CCR2+ cells acquired a characteristic marker of lining macrophages – VSIG4 (**Figure 4C-E**), suggesting that after insult, monocytes can replenish lining macrophages. However, not all CCR2+ cells were VSIG4-positive macrophages (**Figure 4F**), raising further potential avenues for investigating the fate of monocytes recruited by RELM⍺-positive macrophages.

**Figure 4:**
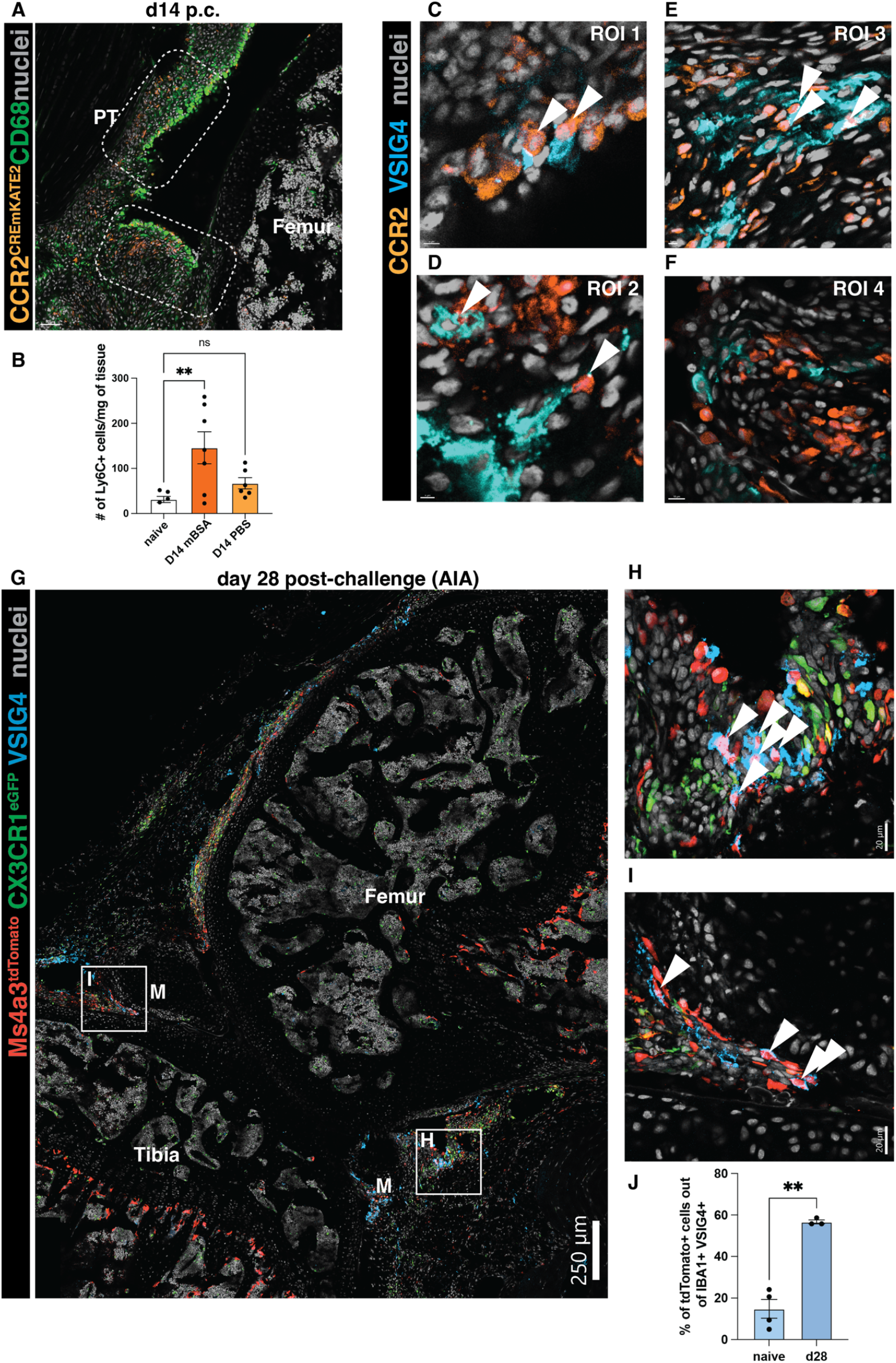
Monocytes can give rise to VSIG4-positive lining macrophages after the resolution of inflammatory arthritis. **(A)** Representative images of CCR2^CreERT-mKATE2^ murine knee sections at the resolution day 14 post-challenge. PT=patellar tendon, M=meniscus. **(B)** Quantification of the number of monocytes at steady state vs at day 14 of AIA in the antigen-injected (mBSA) knee and its contralateral control (PBS). Outliers were identified by the ROUT (Q=1%) test and subsequently removed (n=2); differences were identified by ordinary one-way ANOVA with Dunnett’s multiple comparison test. Each point represents a biological replicate/timepoint. **(C-F)** ROIs from the CCR2^CreERT-mKATE2^ murine knee sections during resolution at day 14 of AIA. Arrows indicate the colocalisation of the CCR2 signal in VSIG4-positive macrophages. **(G)** A representative section from Ms4a3^tdTomato^ CX3CR1^eGFP^ mouse at day 28 of AIA. M=meniscus. **(H-I)** ROIs highlighted in (G); arrows indicate Ms4a3^tdTomato^ VSIG4-double positive cells. **(J)** Quantification of the contribution of monocytes to the lining macrophage pool at steady-state and day 28 of AIA. Each point represents an average of 2-3 positional replicates from 1 biological replicate (n=4 steady state, n=3 d28); Welch’s t-test was used to identify the difference in monocyte contribution between naïve and day 28.

To formally assess the contribution of recruited monocytes into the synovial lining macrophage pool during the resolution of inflammation, we used the Ms4a3^tdTomato^ monocyte tracing model [10], at day 28 post-antigen challenge. Day 28 was chosen instead of day 14, to look at lasting, long-term changes in the synovium post-inflammation. Interestingly, we noted many tdTomato-positive cells in the lining of these mice (**Figure 4G**), many of which were positive for VSIG4 and another classical marker of lining macrophages, CX3CR1 [3, 16, 19] (**Figure 4H, I**). By quantifying the contribution of monocytes to the maintenance of lining macrophages at steady-state and day 28 of AIA, we were able to distinguish that, indeed, at steady-state, only a small proportion of lining macrophages were derived from monocytes [3, 19] (**Figure 4J**). Conversely, at day 28, approximately half of the lining macrophages were derived from monocytes (**Figure 4J**).

Overall, this study identified that monocytes are recruited to the synovial interstitium at the onset of inflammatory arthritis, at least partially by a subset of perivascular CCL2-producing RELM⍺+ macrophages. Monocytes recruited into the synovium can have many functions, including, but not limited to, the replenishment of tissue-resident lining macrophages.

## Discussion

In this brief research report, we identified that RELMα-positive synovial macrophages were located in the perivascular niche, similar to their human counterparts, Lyve1-positive macrophages [4], and functioned as cells sending signals for monocyte recruitment into the interstitium at the onset of inflammatory arthritis, i.e. monocyte chemoattractant CCL2. Finally, we observed that some monocytes appear to be retained in the synovium and even start expressing tell-tale markers of other tissue-resident macrophage subsets, including VSIG4-positive lining macrophages.

Our study clearly demonstrated that synovial interstitial macrophages encompass two distinct subsets, expressing MHCII or RELM⍺, with possibly diverse functions. This is consistent with the comprehensive scRNA-seq profiling of synovial macrophages [3], and different from a recent study, in which expression of MHCII was used to characterise all interstitial macrophages in the synovium [19]. RELMα-positive macrophages are capable producers of CCL2, a main chemoattractant for CCR2-expressing classical monocytes. Similarly, perivascular macrophages in the gut were found to be able to secrete CCL2 in a model of colitis, suggesting that CCL2 production during tissue inflammation may be a shared feature of perivascular macrophage subsets [25].

We previously shown that the influx of monocytes in the antigen-challenged joints was dependent on their expression of CCR2 [24]. Indeed CCR2-positive monocytes were readily recruited into the synovium, at the onset of inflammation. Interestingly, the newly recruited monocytes are specifically localised in the interstitium, unlike neutrophils which accumulate in the lining at the onset of articular inflammation [16]. This phenomenon highlights the dichotomy of synovial tissue-resident macrophage function in the lining and the interstitium, leading to a distinct localisation pattern of myeloid cells at the onset of synovial inflammation.

With progression of inflammation, the recruited monocytes differentiate into macrophages, with differential requirements for the replenishment of interstitial versus lining macrophages at different stages of AIA. It is currently unknown why such specification of the myeloid cell recruitment and localisation exists and how it contributes to arthritis immunopathology, but the finding that lining macrophages are longer-lived than interstitial macrophages [3], and that CCR2-positive cells acquire a tell-tale marker of lining macrophages, VSIG4 [2, 16], during the resolution phase of synovial inflammation, may offer a possible cue. Whether these monocyte-derived lining macrophages have distinct functions from those present prior to the synovial injury is currently unknown, and our report provides a compelling case for further enquiries into this topic beyond the scope of our brief research report.

## Acknowledgements

We thank the Microscopy and Flow Cytometry Facilities at the Kennedy Institute of Rheumatology, namely Dr Kseniya Korobchevskaya, Dr Helena Coker, and Dr Jonathan Webber. Further, we thank Karolina Kaczkowska for colony management. We are grateful to Prof Burkhard Becker (University of Zurich) for providing CCR2^CreER mKate2^ mice.

This work was funded by the Wellcome Trust (Investigator Award 209422/Z/17/Z to I.A. Udalova, K. Zec; PhD Studentship 222344/Z/21/Z to B. Schonfeldova) and the Versus Arthritis (Foundation Fellowship 22568 to K. Zec). The Zeiss LSM 980 was acquired and maintained through grants from the Kennedy Trust for Rheumatology Research.

## Generative AI statement

The authors declare that no Generative AI was used in the creation of this manuscript.

## Data and materials availability

This paper does not report original code or data sets. Any additional information required to reanalyse the data reported in this paper is available from the corresponding authors upon reasonable request.

## Supplementary figure legends

**Supplementary Figure 1:**
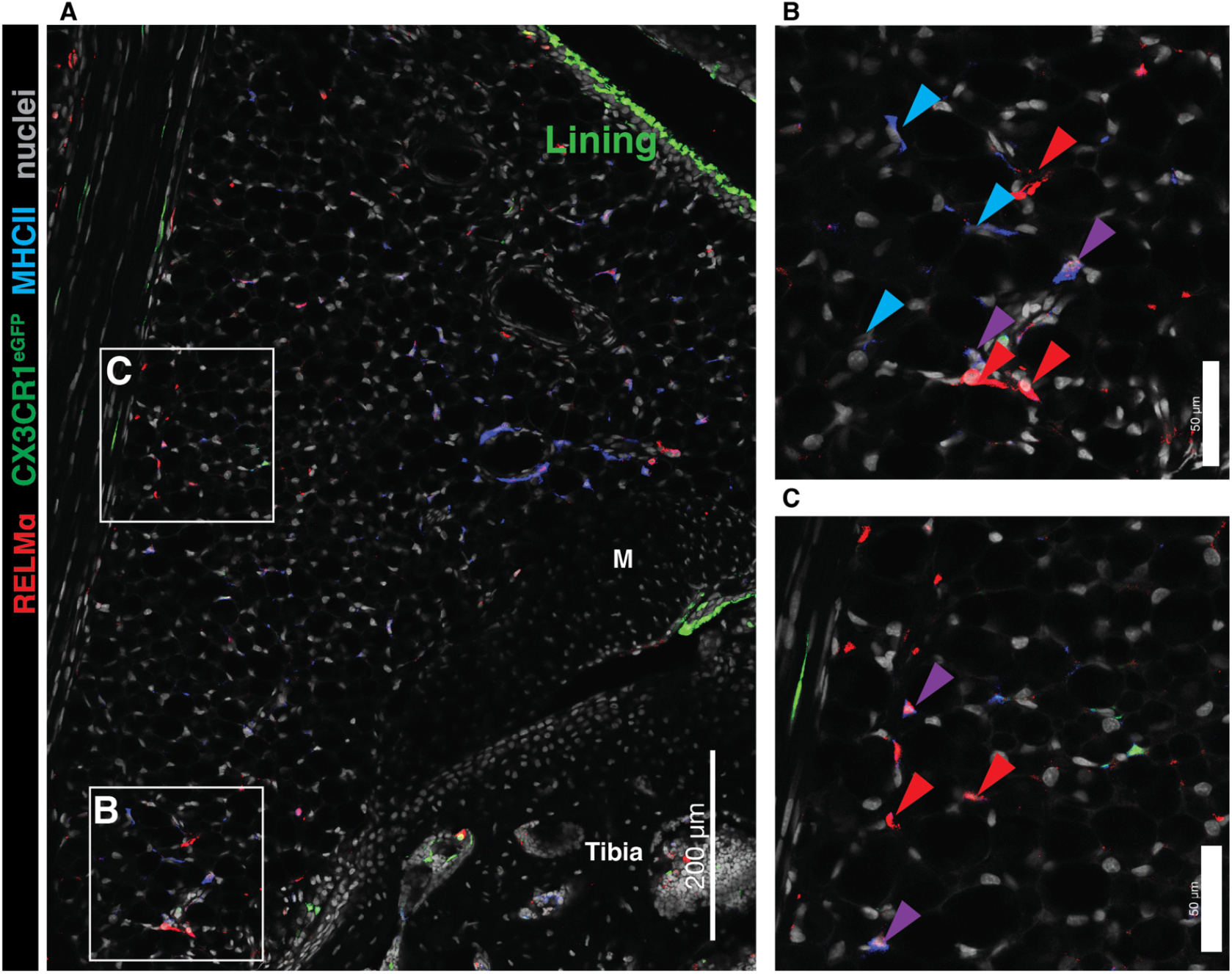
MHCII and RELM⍺ colocalise in some, but can mark different subsets of synovial macrophages. **(A)** A representative image of the naïve synovium, MHCII and RELM⍺ are co-stained and displayed as a pseudochannel with pan-macrophage marker IBA1 to denote macrophages only. The CX3CR1^eGFP^ marks lining macrophages; M=meniscus. **(B-C)** ROIs highlighted in (A), red arrow indicates RELM⍺-positive macrophages, blue arrow marks MHCII-positive macrophages, and purple arrow marks macrophages double-positive for both RELM and MHCII.

**Supplementary Figure 2:**
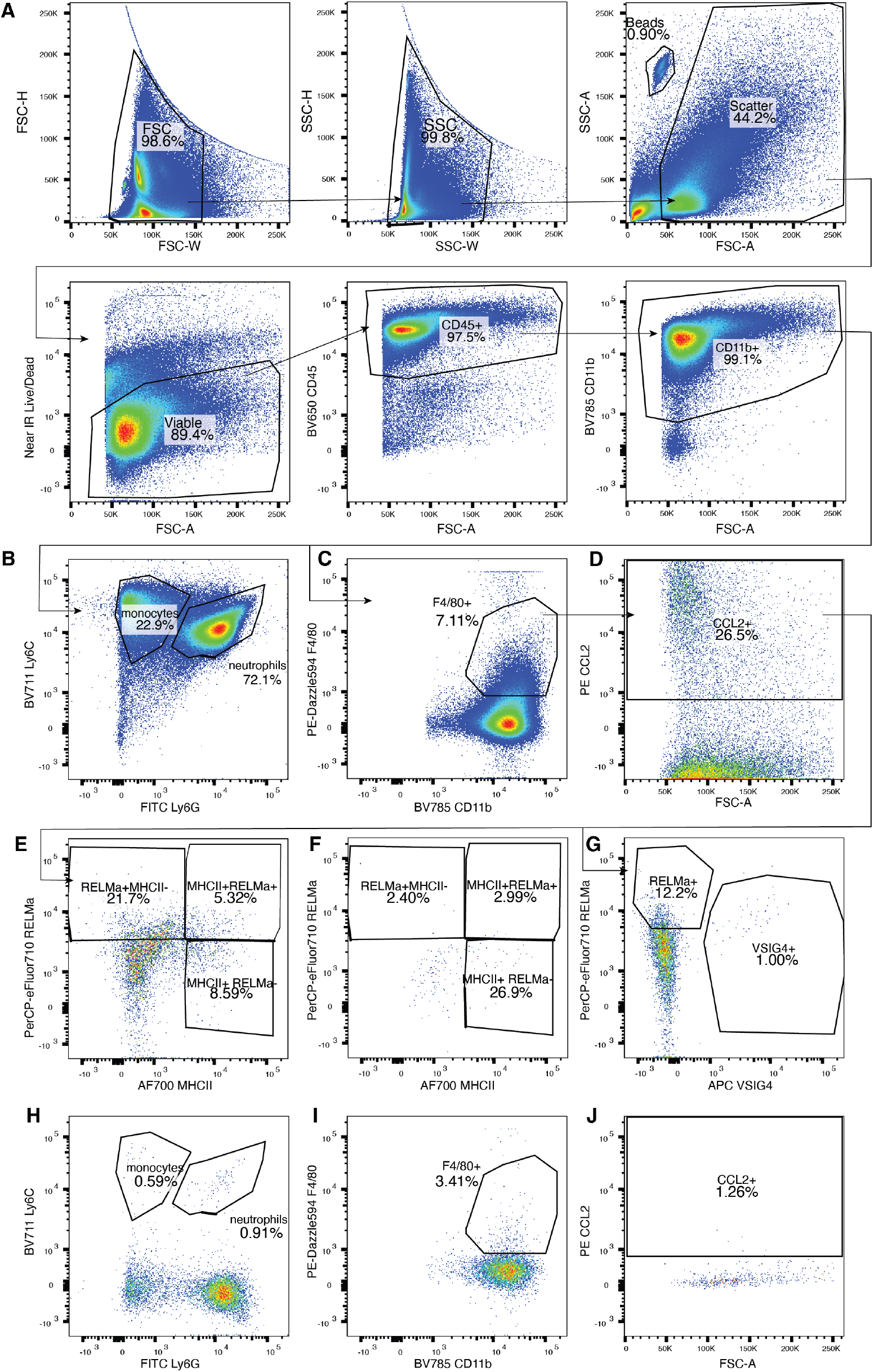
Gating strategy to identify CCL2-producing macrophages at day 1 of AIA. **(A-D)** Identification of the myeloid fraction (A, viable CD45+CD11b+), monocytes (B, viable CD11b+Ly6C+Ly6G-), neutrophils (B, viable CD11b+Ly6C+Ly6G+), macrophages (C, viable CD11b+Ly6C-Ly6G-F4/80+), and CCL2-producing macrophages (D, viable CD11b+Ly6C-Ly6G-F4/80+CCL2+). **(E-F)** Gating for RELMα+, MHCII+, and RELMα+MHCII+ with FMO control for RELMα (F). **(G)** Identification of VSIG4+ macrophages. **(H-J)** FMO controls for Ly6C (H), F4/80 (I), and CCL2 (J) staining.

